# Stromal KITL/SCF promotes pancreas tissue homeostasis and restrains tumor progression

**DOI:** 10.1101/2024.07.29.605485

**Authors:** M. Kathrina Oñate, Chet Oon, Sohinee Bhattacharyya, Vivien Low, Canping Chen, Xiaofan Zhao, Ziqiao Yan, Yan Hang, Seung K. Kim, Zheng Xia, Mara H. Sherman

## Abstract

Components of normal tissue architecture serve as barriers to tumor progression. Inflammatory and wound-healing programs are requisite features of solid tumorigenesis, wherein alterations to immune and non-immune stromal elements enable loss of homeostasis during tumor evolution. The precise mechanisms by which normal stromal cell states limit tissue plasticity and tumorigenesis, and which are lost during tumor progression, remain largely unknown. Here we show that healthy pancreatic mesenchyme expresses the paracrine signaling molecule KITL, also known as stem cell factor, and identify loss of stromal KITL during tumorigenesis as tumor-promoting. Genetic inhibition of mesenchymal KITL in the contexts of homeostasis, injury, and cancer together indicate a role for KITL signaling in maintenance of pancreas tissue architecture, such that loss of the stromal KITL pool increased tumor growth and reduced survival of tumor-bearing mice. Together, these findings implicate loss of mesenchymal KITL as a mechanism for establishing a tumor-permissive microenvironment.

**Statement of significance:** By analyzing transcriptional programs in healthy and tumor-associated pancreatic mesenchyme, we find that a sub-population of mesenchymal cells in healthy pancreas tissue express the paracrine signaling factor KITL. Loss of mesenchymal KITL is an accompanying and permissive feature of pancreas tumor evolution, with potential implications for cancer interception.

## Introduction

Though evidence that normal tissue components can restrain tumor progression dates back to the 1960s (1), the specific tissue-level barriers to plasticity and tumor outgrowth remain largely unknown. Mechanisms maintaining tissue homeostasis and limiting tumorigenesis include epithelial-epithelial interactions, such as regenerative or competitive epithelial functions (2–4); epithelial-immune interactions, wherein innate (5, 6) or adaptive (7, 8) immune cells clear mutant cells or early pre-invasive lesions; and epithelial-mesenchymal interactions, with evidence that mesenchymal elements such as normal tissue fibroblasts can restrain growth of transformed epithelial cells (9–11). These mechanisms co-exist with, and likely interact functionally with, epithelial cell-intrinsic tumor suppressor gene products, together creating genetic, cellular, and tissue-level checks on cancer development. While epithelial cell-intrinsic mechanisms of tumor suppression have been studied extensively, and we have advanced considerably in our understanding of anti-tumor functions of the immune systems, mechanisms underlying the tumor-restraining potential of normal mesenchyme largely have not been identified.

Fibro-inflammatory reactions create tissue contexts permissive to tumor progression (12, 13). Local or systemic cues, including paracrine signaling from transformed epithelial cells or diverse sources of tissue damage, cause alterations to resident mesenchymal cells such as transition from quiescent fibroblasts to activated myofibroblasts and changes to or accumulation of immune cells. This wound healing reaction helps to promote plasticity in the epithelial compartment and overcome intrinsic barriers to tumor formation and growth (14). Consistent with this notion, though normal primary fibroblasts can suppress hyperplastic growth of mammary epithelial cells *in vivo*, this outgrowth is supported by activated, myofibroblastic stroma (15, 16). Though the causal link between inflammation and cancer has been appreciated for some time (17), recent studies of patient tissues have begun to identify specific mechanisms by which inflammatory insults promote cancer development. For example, environmental pollutants result in an accumulation of IL-1β-producing macrophages in the lung, and this inflammatory signaling drives plasticity in the lung epithelium to promote tumorigenesis (18). Further study of the specific signals engaged by healthy or inflamed tissue components to restrain or promote tumorigenesis, respectively, may point to new avenues for early cancer intervention.

The recent discovery that normal, adult human pancreas tissue harbors up to hundreds of *KRAS*-mutant pre-invasive lesions impels the field to understand intracellular and heterocellular mechanisms restraining neoplastic progression in the pancreas (19, 20). To assess a role for mesenchymal cell state alterations in the transition from a homeostatic to tumor-permissive tissue context, we performed transcriptional profiling of healthy and cancer-associated pancreatic mesenchyme using an established fate mapping mouse model (21). These experiments focused on pancreatic stellate cells (PSCs), tissue-resident mesenchymal cells which serve as cells of origin for PDAC CAFs. We found that this mesenchymal lineage in normal human and murine pancreas tissue expresses KITL—this lineage has lipid storage capacity and co-expresses the leptin receptor (LEPR), with parallels to LEPR-positive mesenchyme previously implicated in tissue homeostasis in the bone marrow (22) and brown adipose tissue (23). Mesenchymal KITL expression is lost during tumor evolution and acquisition of a cancer-associated fibroblast (CAF) stromal phenotype, with functional significance for tissue state and tumorigenic potential.

## Results

To assess stromal evolution during stepwise tumorigenesis, we applied a previously established fate mapping approach (21) to analyze the contributions PSCs to the stroma of normal pancreas tissue, pancreatic intraepithelial neoplasia (PanINs), and invasive PDAC. To this end, we generated a dual recombinase genetically engineered mouse model of the genotype *Kras^FSF-G12D/+^;Trp53^FRT/+^;Pdx1-FlpO;Rosa26^mTmG/+^;Fabp4-Cre* (Figure 1A) and assessed the presence of GFP^+^ stroma, indicating a lipid-storing origin. While GFP^+^ PSCs were found in normal pancreas tissue as expected, very few were positive for PDPN, a cell surface marker upregulated upon fibroblast activation in PDAC. We found GFP^+^PDPN^+^ cells associated with low-grade PanIN lesions as well as invasive cancer in this model (Figure 1B & 1C), with a significant increase in PSC-derived fibroblastic cells in the context of PDAC compared to pre-invasive lesions. In normal pancreas tissue and in tumors, PSCs or PSC-derived CAFs had a spatial distribution similar to the reported tissue distribution of stellate cells in the liver, the other tissue in the body where these mesenchymal cells reside. Hepatic stellate cells (HSCs) are found in perivascular regions in close proximity to endothelial cells, and adjacent to neighboring parenchymal cells (24). We found PSCs in normal pancreas tissue similarly to localize in perivascular regions, and in the tissue parenchyma spatially poised for cell-cell communication with epithelial cells (Figure 1D). This spatial distribution was conserved upon differentiation to a CAF phenotype, as GFP^+^ CAFs were found both immediately adjacent to and distant from endothelial cells in the genetically engineered PDAC model (Figure 1E). Similar results were observed in an orthotopic model wherein PDAC cells derived from the *Kras^LSL-G12D/+^;Trp53^LSL-R172H/+^;Pdx1-Cre* (KPC) autochthonous model were implanted into syngeneic *Rosa26^mTmG/+^;Fabp4-Cre* hosts (Figure 1F). These observations indicate that PSCs contribute to the stromal microenvironment throughout pancreatic tumorigenesis and are spatially distributed to engage in direct cell-cell contact with both endothelial cells and epithelial cells in healthy and cancerous pancreas tissue.

**Figure 1:**
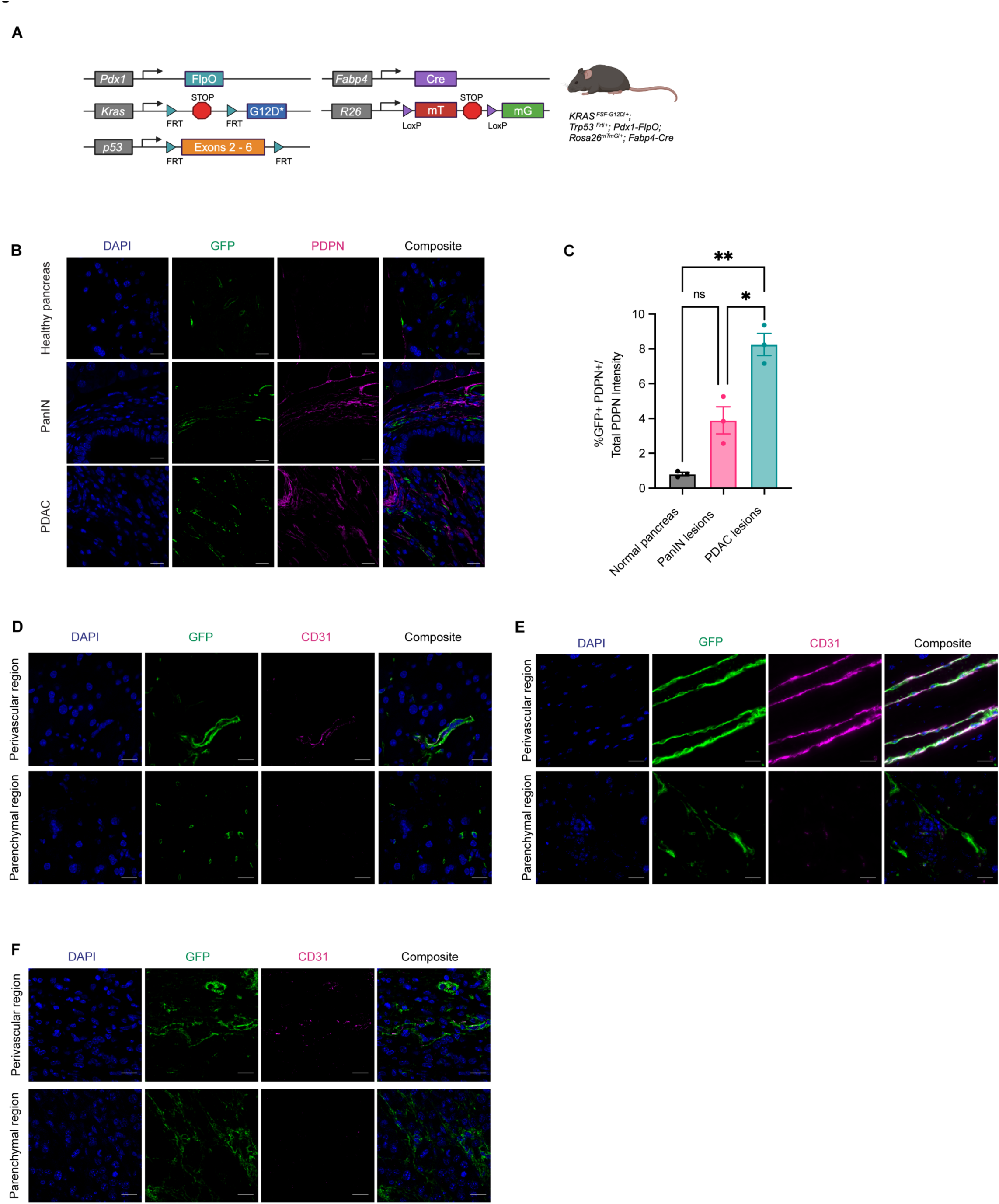
Pancreatic stellate cells contribute to the stromal microenvironment throughout tumorigenesis. **A,** Genetic schema of *Kras^FSF-G12D/+^;Trp53^FRT/+^;Pdx1-FlpO;Rosa26^mTmG/+^;Fabp4-Cre* murine model. **B,** Representative images of IHC staining for GFP (green) and Podoplanin (PDPN, magenta) among normal pancreas, PanIN lesions, and mPDAC lesions. Scale bar, 10 μm. **C,** IHC staining quantification of percent GFP^+^PDPN^+^ (double-positive) cells over total PDPN^+^ expression among the 3 disease states in **B** (n = 3). **D,** Representative images of IHC staining for GFP (green) and CD31 (magenta) within normal pancreas (n = 5). Scale bar, 10 μm. **E,** Representative images of IHC staining for GFP (green) and CD31 (magenta) within GEMM pancreata (n = 3). Scale bar, 20 μm. **F,** Representative images of IHC staining for GFP (green) and CD31 (magenta) within pancreata of KPC-derived orthotopically implanted PDAC in *Rosa26^mTmG/+^;Fabp4-Cre* mice (n = 3). Scale bar, 10 μm.

We next assessed alterations in expression of cell surface ligands or receptors in this mesenchymal lineage during pancreatic tumorigenesis. We reasoned that paracrine signaling factors important for normal tissue architecture may be lost from the mesenchyme during the transition from normal tissue homeostasis to cancer. To identify candidate paracrine factors associated with normal mesenchymal function whose loss may be tumor-permissive, we analyzed the transcriptional profiles of PSCs and PSC-derived CAFs. To this end, we sorted GFP^+^ PSCs and GFP^+^PDPN^+^ CAFs from healthy pancreas tissue and PDAC, respectively, and performed single-cell RNA-seq (scRNA-seq) to assess gene expression differences in this cellular compartment within and across tissue states. As expected, these cells in normal pancreas and PDAC were pervasively positive for mesenchymal marker *Vim* and almost all positive for pan-tissue fibroblast markers *Pi16* or *Col15a1* (25) (Figure 2A). Though not all cells expressed one of these two universal fibroblast markers, we note that PSCs are not strictly fibroblasts albeit fibroblast-like. While PSCs and PSC-derived CAFs are partially perivascular as described above, these cells lacked expression of classical pericyte markers such as *Cspg4* (encoding NG2) and *Rgs5* (Supplementary Figure S1A). Interestingly, the sub-population in normal pancreas tissue lacking universal fibroblast markers expressed both *Vim* and genes generally associated with a macrophage identity, such as *Csf1r* and *Adgre1* (encoding F4/80) (Supplementary Figure S1B). However, when we stained for GFP and macrophages in pancreas tissues we detected no overlap (Supplementary Figure S1C), suggesting that this sub-population of cells in normal pancreas tissue may be fibrocyte-like or otherwise express some macrophage-associated genes without assuming a macrophage identity. In the context of cancer, PSCs gained expression of immune-modulatory cytokines such as *il6* and *Il33* as expected for CAFs (26) and pervasively expressed extracellular matrix (ECM) components such as *Col1a2* (Figure 2B). These results validate activation of PSCs to a CAF phenotype in PDAC.

**Figure 2:**
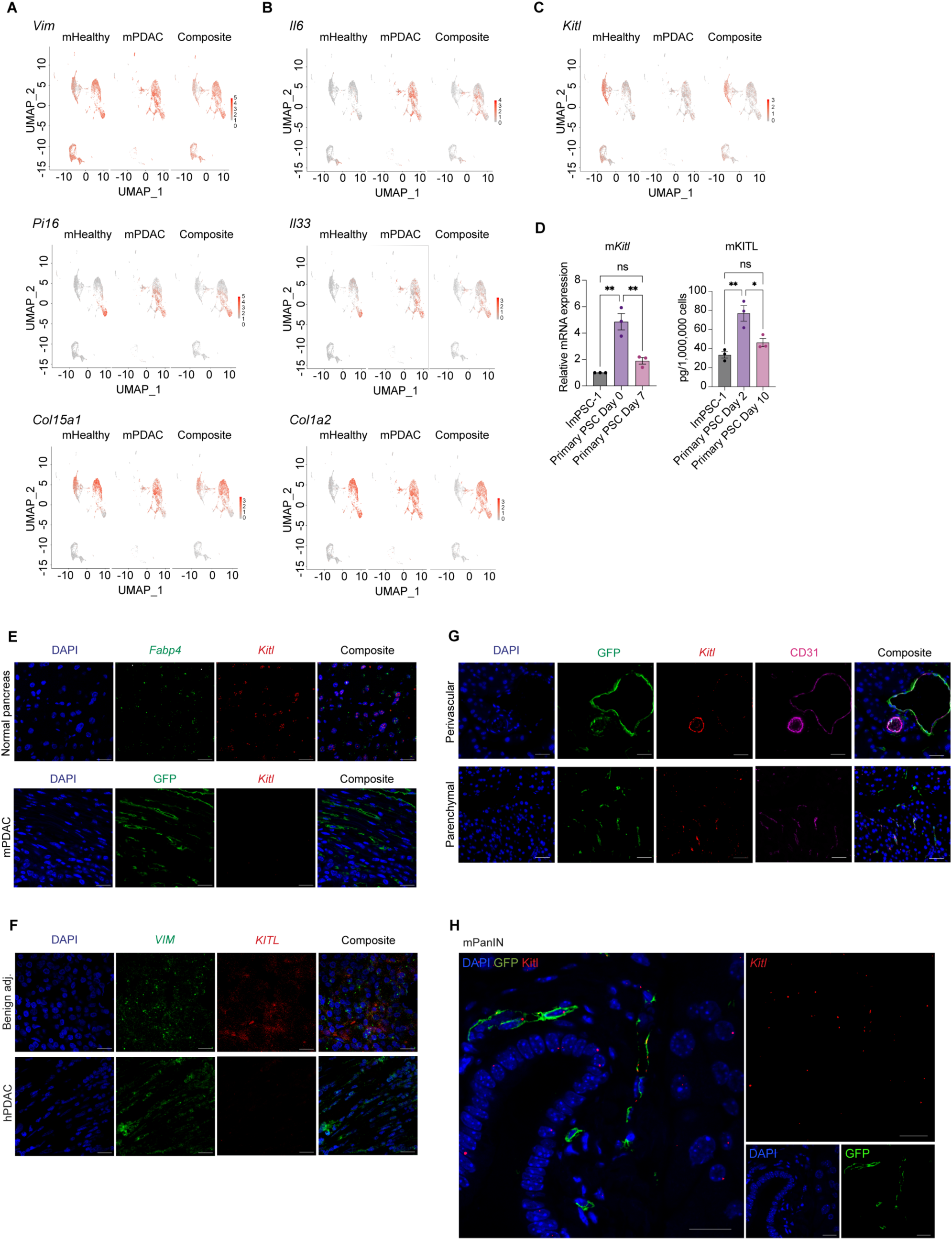
Mesenchymal KITL loss within PSCs accompanies pancreatic tumorigenesis. **A,** UMAP visualization of *Vim*, *Pi16*, and *Col15a1* gene expression in normal pancreatic stellate cells (PSCs) and PSC-derived cancer associated fibroblasts scRNA-seq dataset (n = 2 replicates pooled from n = 5 mice per arm). **B,** UMAP visualization of *Il6*, *Il33*, and *Col1a2* gene expression from normal pancreatic stellate cells (PSCs) and PSC-derived cancer associated fibroblasts scRNA-seq dataset (n = 2 replicates pooled from n = 5 mice per arm). **C,** UMAP visualization of *Kitl* transcript expression in normal pancreatic stellate cells (PSCs) and PSC-derived cancer associated fibroblasts scRNA-seq dataset (n = 2 replicates pooled from n = 5 mice per arm). **D,** Left: qRT-PCR analysis of *Kitl* in quiescent (Day 0) and activated (Day 7) primary pancreatic stellate cells (PSCs). Right: Quantikine ELISA KITL measurement of supernatant collected from primary PSCs in pre-activated (Day 2) and activated state (Day 10) after 48 hours incubation with media change on Day 8 to harvest for Day 10 sample. Immortalized ImPSC-1 included as reference point. Data represents biological triplicate plotted as mean ± SEM. Significance was determined by ordinary one-way ANOVA; ns = not significant, *P≤ 0.05, **P≤ 0.01. **E,** Representative images of RNA FISH staining for *Fabp4* (green) and *Kitl* (red) in murine normal pancreas (n = 3). Scale bar, 10 μm. Below, representative RNAScope staining of GFP (green) protein and *Kitl* (red) mRNA in PDAC from the GEMM depicted in **1A** (n = 3). Scale bar, 10 μm. **F,** Representative images of RNA FISH staining for *VIM* (green) and *KITL* (red) in human PDAC tissues between benign adjacent and PDAC regions (n = 3). Scale bar, 10 μm. **G,** Representative images of RNAScope staining for *Kitl* (red) mRNA expression, GFP (green) and CD31 (magenta) in murine normal pancreas from *Rosa26^mTmG/+^;Fabp4-Cre* mice (n = 3). Scale bar, 20 μm. **H,** Representative images of RNAScope staining for GFP (green) protein and *Kitl* (red) mRNA in GEMM low-grade PanIN (n = 3). Scale bar, 20 μm.

We next focused on paracrine signaling factors expressed in healthy pancreatic mesenchyme and lost in PDAC which may represent barriers to tumor progression. We noted expression of *Kitl* (also known as stem cell factor or SCF) in normal pancreas tissue but lost in CAFs (Figure 2C), supported by pseudo-time analysis (Supplementary Figure S2A). KITL expression has not previously been reported in normal pancreas tissue, and was of interest to us in light of the significance of KITL-positive mesenchyme in the perivascular niche of the bone marrow, where stromal KITL is crucial for normal tissue structure and function (22). Further, HSCs in the developing liver are critical sources of KITL to support the hematopoietic stem cell niche (27), providing precedent for functionally significant KITL production by stellate cells. KITL-positive mesenchyme in the bone marrow express the leptin receptor (LEPR), and we detected low levels of *Lepr* expression among normal PSCs by scRNA-seq (Supplementary Figure S2B). We validated these findings by isolating primary PSCs from healthy pancreas tissue and activating them to a CAF phenotype in culture: These cells expressed *Kitl* and *Lepr* in their normal tissue state but progressively lost expression of both factors upon activation to a CAF-like state (Figure 2D, Supplementary Figure S2C & S2D). We next validated expression of *Kitl* in intact murine pancreas tissue. By RNA *in situ* hybridization (ISH, due to lack of specific antibodies, using branched cDNA hybridization), we detected *Kitl* expression in mesenchymal cells of normal pancreas tissue which share markers with PSCs, while *Kitl* expression was lost among CAFs in PDAC (Figure 2E). We also combined RNA ISH for *Kitl* (here using RNAscope, compatible with protein co-staining) with immunohistochemistry (IHC) for GFP on pancreas tissue from *Rosa26^mTmG/+^;Fabp4-Cre* mice and confirmed *Kitl* expression in fate-mapped PSCs. Specificity of our *Kitl* probe was confirmed by reduction in mesenchymal *Kitl* signal in pancreas tissues from *Kitl^flox/flox^;Fabp4-Cre* mice (Supplementary Figure S2E). We extended these analyses to human pancreas tissue, and performed RNA ISH for *KITL* and mesenchymal marker *VIM*. While benign human pancreas harbored *KITL*-positive mesenchyme, CAFs within human PDAC lost *KITL* expression, consistent with observations in mice (Figure 2F). As perivascular mesenchyme is a critical source of KITL in other tissues (22, 27), we assessed the spatial distribution of mesenchymal *Kitl* expression by combining *Kitl* RNA ISH with IHC for CD31 and GFP on pancreas tissues from *Rosa26^mTmG/+^;Fabp4-Cre* mice. We found that PSCs express *Kitl* in both perivascular regions and when not adjacent to endothelial cells (Figure 2G), suggested that KITL from PSCs is poised to signal to multiple neighboring cell types. To assess the stage of pancreatic tumorigenesis at which mesenchymal *Kitl* expression is lost, we combined RNA ISH for *Kitl* and IHC for GFP (to indicate PSCs and PSC-derived CAFs) on tissues from *Kras^FSF-G12D/+^;Trp53^FRT/+^;Pdx1-FlpO;Rosa26^mTmG/+^;Fabp4-Cre* mice and noted retention of *Kitl* expression among GFP-positive stromal cells associated with low-grade PanIN lesions identified by a pathologist (Figure 2H), suggesting that loss of stromal *Kitl* accompanies late stages of pancreatic tumorigenesis. Expression of *Kitl* by some GFP-negative cells was noted within these areas of low-grade PanIN as well. Together, these analyses revealed expression of *Kitl* by a lineage of healthy pancreatic mesenchyme in mice and humans which is lost upon transition to a CAF phenotype in invasive cancer.

We next addressed the functional significance of KITL in pancreatic mesenchyme, and assessed the consequence of stromal KITL loss for tissue homeostasis. First, we questioned the cell-intrinsic impact of KITL signaling on pancreatic mesenchymal cells. To address this, we generated loss- and gain-of-function systems in cell culture by knocking down or overexpressing *Kitl* in immortalized PSCs (28) using shRNA or introduction of the *Kitl* ORF, respectively (Supplementary Figure S3A). PSCs in culture express low but detectable levels of *Kitl* (Figure 2D), so we reasoned that gene expression changes observed with *Kitl* overexpression would reflect downstream transcriptional programs in healthy mesenchyme while *Kitl* knockdown would reflect consequences of gene expression changes upon transition to a CAF state. In culture, PSCs express low but detectable levels of *Kit* (encoding c-KIT) (Supplementary Figure S3B), the paracrine signaling partner for KITL, such that PSC monoculture seemed a reasonable *in vitro* model to begin assessing how KITL signaling impacts pancreatic mesenchyme. To this end, we analyzed the transcriptional profiles of *Kitl*-knockdown and *Kitl*-overexpressing PSCs, together with appropriate controls, by RNA-seq. We prioritized gene expression changes resulting from *Kitl* overexpression as this cell line is activated and therefore CAF-like, though *Kitl* knockdown was indeed achievable. Restoring *Kitl* expression caused upregulation of genes involved in cell adhesion and extracellular matrix or collagen organization, including integrins, laminins, cadherins, and protocadherins (Figure 3A & 3B), suggesting potential involvement of stromal KITL in regulation of normal tissue architecture. Conversely, *Kitl* knockdown led to upregulation of genes involved in inflammatory processes, including genes involved in complement or interferon signaling, together with downregulation of cell adhesion genes (though many of the genes positively regulated by *Kitl* signaling were expressed at a very low level in control cells and were not significantly downregulated further upon *Kitl* knockdown) (Supplementary Figure S3C & S3D). These results suggested that mesenchymal KITL may promote pancreas tissue homeostasis, prompting us to move into *in vivo* modeling of KITL function.

**Figure 3:**
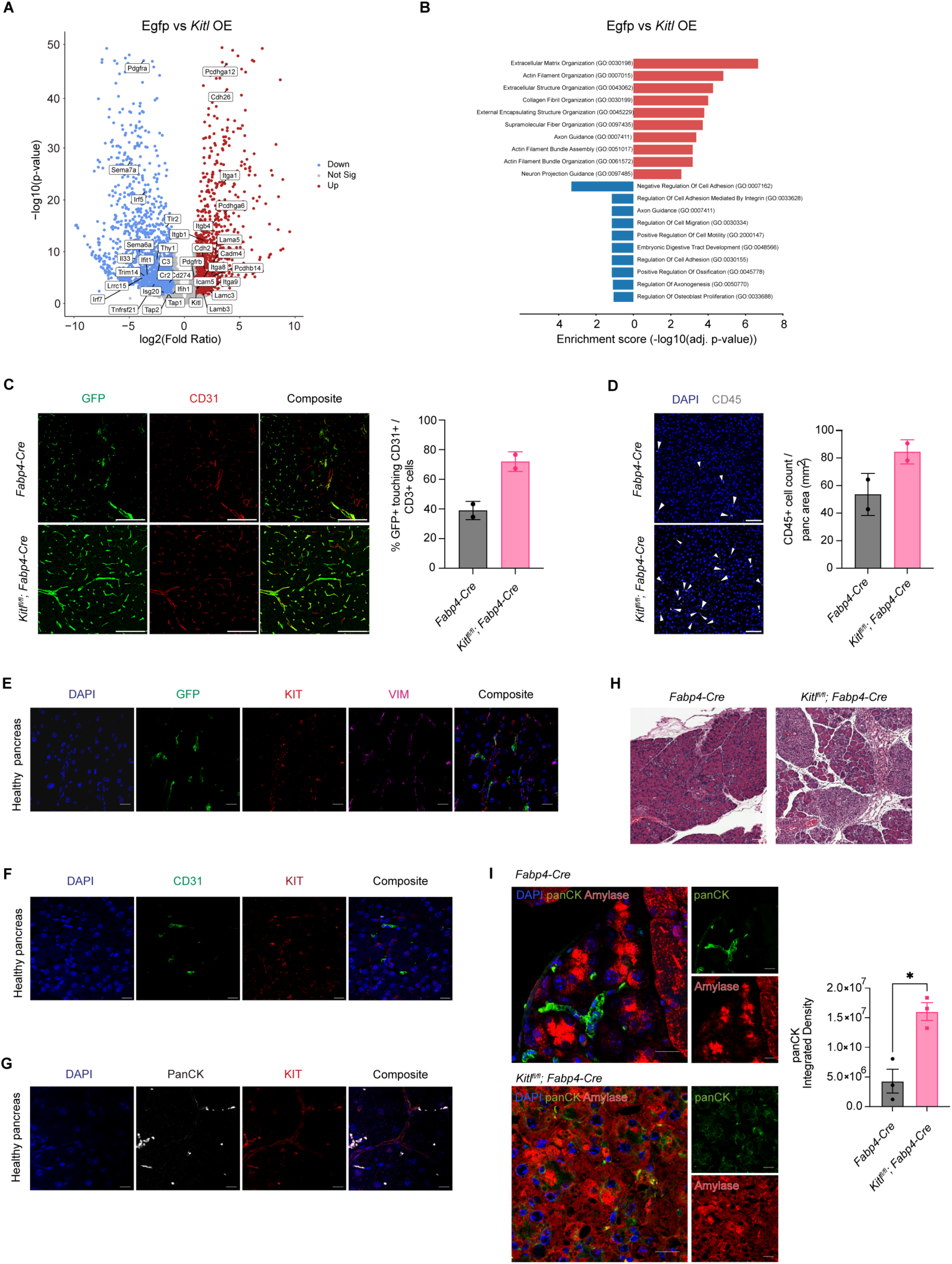
KITL regulates PSC state and pancreas tissue homeostasis. **A,** Volcano plot of all upregulated, non-significant, and downregulated differentially expressed genes (DEGs) as defined by the Wald test (p.adj <0.05 and log_2_FC >1) from *Kitl* overexpression (*Kitl* OE) ImPSC-1 bulk-RNA seq dataset with representative gene labels included. Data represent 3 biological repeats. **B,** Gene ontology (GO) analysis of upregulated and downregulated genes in immortalized pancreatic stellate cells (ImPSC-1) overexpressing *Kitl*. Top 10 enrichment categories ranked by adjusted p-values plotted in each direction. **C,** Representative images of CODEX staining (left) and quantification (right) for GFP (green) and CD31 (red) in normal pancreas from *Fabp4-Cre* or *Kitl^fl/fl^;Fabp4-Cre* mouse model (n= 2 mice per arm). Scale Bar: 100 μm. Data are represented as mean ± SD. **D,** Representative images of CODEX composite staining (left) and quantification (right) for CD45 (white) in normal pancreas from *Fabp4-Cre* or *Kitl^fl/fl^;Fabp4-Cre* mouse model (n= 2 mice per arm). Scale Bar: 50 μm. Data are represented as mean ± SD. **E,** Representative images of IHC staining for GFP (green), cKIT receptor (KIT, red), and VIM (magenta) in healthy murine pancreas from *Rosa26^mTmG/+^;Fabp4-Cre* mice. Scale bar, 10 μm. **F,** Representative images of IHC staining for CD31 (green) and cKit receptor (KIT, red) in healthy murine pancreas. Scale bar, 10 μm. **G**, Representative images of IHC staining for panCK (white) and cKit receptor (KIT, red) in healthy murine pancreas. Scale bar, 10 μm. **H**, Representative H&E images between caerulein-treated *Fabp4-Cre* and *Kitl^fl/fl^;Fabp4-Cre* mice. Scale bar, 100 μm. **I**, Representative images of IHC staining (left) for panCK (green) and Amylase (red) between caerulein-treated *Fabp4-Cre* and *Kitl^fl/fl^;Fabp4-Cre* mice. Scale bar, 10 μm. PanCK quantification on right (n = 3 mice per arm). For comparisons between two groups, Student’s two-tailed t-test was used. Data are represented as mean ± SEM. *P < 0.05.

To assess the relevance of mesenchymal KITL signaling for pancreas tissue architecture, we analyzed the consequences of conditional *Kitl* loss using *Kitl^flox/flox^;Fabp4-Cre* mice compared to *Fabp4-Cre* controls in the settings of homeostasis and tissue injury. First, we analyzed these tissues under normal, homeostatic conditions, and crossed in a *Rosa26^mTmG/+^* allele to enable visualization of PSCs based on GFP expression in these tissues. Based on our transcriptional profiling results, we compared tissue microenvironments in *Rosa26^mTmG/+^;Kitl^flox/flox^;Fabp4-Cre* mice compared to *Rosa26^mTmG/+^;Fabp4-Cre* controls using co-detection by indexing (CODEX), a barcode-based, multiplexed imaging approach (29). While total VIM-positive and CD31-positive cell abundance was not different between genotypes (Supplementary Figure S4A & S4B), we observed clear changes to the perivascular niche with loss of mesenchymal *Kitl* including an increase in GFP-positive mesenchyme adjacent to endothelial cells (Figure 3C). We also observed an increase in CD45-positive leukocytes within normal pancreas tissue when stromal *Kitl* was perturbed (Figure 3D). We also noted a trend towards decreased α-SMA-positive, VIM-positive cells with *Kitl* perturbation (Supplementary Figure S4C)—as fibroblasts are α-SMA-negative in normal pancreas tissue, this likely reflects a reduction in contractility of vascular smooth muscle cells. To assess potentially cellular receivers of mesenchymal KITL which participate in paracrine signaling, we stained pancreas tissues from *Rosa26^mTmG/+^;Fabp4-Cre* mice for GFP, VIM, and KITL receptor KIT. We found that KIT-positive cells were found adjacent to GFP-positive mesenchyme, consistent with the potential for cell-cell communication (Figure 3E). As PSCs are localized in perivascular regions as well as next to pancreatic epithelium, but KIT-positive cells were few in number in pancreas tissue, we reasoned that acinar cells were unlikely to be the cellular source of KIT but that CD31-positive endothelial cells and cytokeratin-high ductal epithelial cells may be relevant KIT-positive cell populations. Consistent with this notion, IHC demonstrated KIT expression by sub-populations of ductal epithelial cells and few endothelial cells (Figure 3F & 3G). To confirm these results, we analyzed KIT expression by flow cytometry with co-stains for CD45 (immune cells), CD31 (endothelial cells), or EpCAM (epithelial cells), reasoning that KIT-positive cells negative for these three additional markers represent KIT-positive mesenchyme. KIT-positive cells were found in the EpCAM-positive fraction, consistent with a ductal epithelial identity, and were rarely but measurably positive for CD31 or CD45 (Supplementary Figure S4D & S4E), consistent with our IHC. We also noted a KIT-expressing population negative for these markers, which may be a population of mesenchymal cells expressing KIT at too low a level for detection by IHC. We also note that the fairly high proportion of KIT-positive cells among live cells in our flow cytometry experiments likely reflects substantial acinar cell death during preparation of single cell suspensions, as acinar cells appear to be negative for KIT and we have likely therefore enriched for KIT-positive cells. In light of measurable albeit modest differences to tissue structure upon loss of mesenchymal *Kitl*, we assessed the consequences of this KITL pool in the setting of tissue damage. For this, we subjected *Kitl^flox/flox^;Fabp4-Cre* mice and *Fabp4-Cre* controls to acute pancreatitis by administering repeated injections of the cholecystokinin analog caerulein, or saline as a vehicle control. As expected, in control mice, caerulein induced a mild inflammation characterized by edema and leukocyte accumulation evident by hematoxylin and eosin staining (Figure 3H). However, in *Kitl* conditional knockout mice, caerulein led to far more pronounced tissue inflammation, as well as greater alterations to the epithelial compartment which we speculated may represent metaplasia or altered epithelial plasticity. To assess this, we co-stained tissues from caerulein-treated mice with amylase (acinar cell marker) and pan-cytokeratin (ductal cell marker), which indicated an increase in ductal marker expression in the inflamed *Kitl* conditional knockout mice (Figure 3I) along with evidence of amylase/pan-cytokeratin co-staining of individual cells. Together, these results implicate mesenchymal KITL in regulation of pancreas tissue homeostasis such that KITL downregulation promotes inflammation and perturbation of normal tissue architecture.

We next addressed the potential of stromal KITL to regulate pancreatic tumor growth. We performed orthotopic implantation of KPC-derived PDAC cells from a pure C57BL/6J background into syngeneic *Kitl^flox/flox^;Fabp4-Cre* mice or *Fabp4-Cre* controls. Despite the aggressive nature of this mouse model, we found that loss of mesenchymal *Kitl* significantly accelerated tumor growth (Figure 4A) and increased tumor weights and tumor burden at experimental endpoint (Figure 4B & 4C). We then repeated these experiments using moribundity as an experimental endpoint instead of a fixed timepoint. Consistent with the tumor growth measurements, survival studies revealed that loss of mesenchymal *Kitl* significantly shortened survival compared to mice in KITL-expressing hosts (Figure 4D). We characterized the mesenchymal compartment of these tumors by staining for PDPN (pan-CAF marker) and α-SMA (myofibroblast-like CAF marker) and found similar CAF abundance in tumors across genotypes (Figure 4E), consistent with the notion that mesenchymal KITL regulates tissue homeostasis but is lost in an established tumor microenvironment. As implantable models involve introduction of cells which have already undergone malignant transformation into pancreas tissue, these results suggest that mesenchymal KITL expression represents a tissue barrier to PDAC progression at least in part independent of epithelial cell-intrinsic tumor suppression mechanisms.

**Figure 4:**
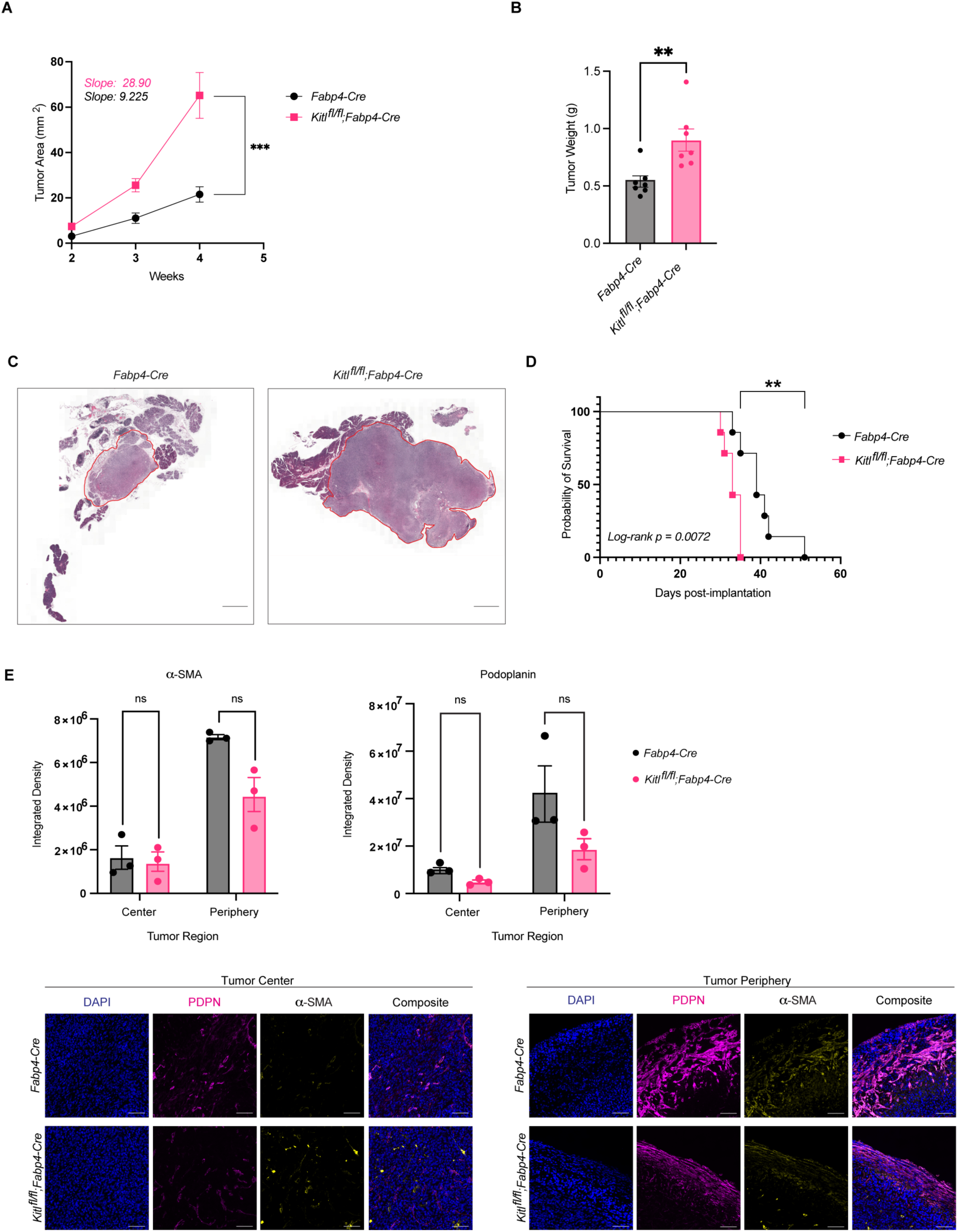
Mesenchymal KITL restrains pancreatic tumor growth. **A,** Average tumor area (mm^2^) between *Fabp4-Cre* control and *Kitl^fl/fl^;Fabp4-Cre* mice, injected with KPC-derived murine PDAC cells 6419c5 (n = 7 mice per arm). Data are represented as mean ± SEM. Slopes tabulated via simple linear regression analysis. **B,** Tumor weights (g) at experimental endpoint between *Fabp4-Cre* control and *Kitl^fl/fl^;Fabp4-Cre* mice, injected with KPC-derived murine PDAC cells 6419c5 (n = 7 mice per arm). Data are represented as mean ± SEM. **C,** Representative H&E images of *Fabp4-Cre* control and *Kitl^fl/fl^;Fabp4-Cre* mice injected with KPC-derived murine PDAC cells 6419c5, at the same experimental endpoint. Scale bar, 1mm. **D,** Kaplan–Meier plot depicting percent probability of survival between *Fabp4-Cre* control and *Kitl^fl/fl^;Fabp4-Cre* mice, injected with KPC-derived murine PDAC cells 6419c5 (n = 7 mice per arm). Log-rank p value = 0.0072. **E,** Representative images of IHC staining (bottom) and quantification (top) of αSMA and podoplanin (PDPN) between *Fabp4-Cre* control mice and *Kitl^fl/fl^;Fabp4-Cre* mice, injected with KPC-derived murine PDAC cells 6419c5 (n = 3 mice per arm). Data are represented as mean ± SEM. *, P < 0.0332; **, P < 0.0021; ***, P < 0.0002; ****, P < 0.0001 Mann-Whitney unpaired t test; ns, not significant.

## Discussion

In this study, we provide evidence that a cell population in normal pancreatic mesenchyme expresses KITL/SCF; that stromal downregulation of KITL is an accompanying feature of pancreatic tumorigenesis, as CAFs derived from these KITL-positive cells retain a lineage label but do not retain KITL expression; and that, functionally, stromal KITL is a barrier to tumor progression in pancreas tissue. The recent reports of abundant, *KRAS*-mutant, pre-invasive lesions throughout examined cohorts of PDAC-free human pancreas tissues (19, 20) compared to the relatively low frequency of PDAC across the general population indicates the pervasive relevance of tumor suppression mechanisms in the adult pancreas. These mechanisms likely include epithelial cell-intrinsic mechanisms promoting, among other things, genome stability and susceptibility to immune surveillance; functions of the immune system, potentially including clearance of highly mutated epithelial cells with tumorigenic potential; and functions of the non-immune stroma. Within the non-immune stroma, mesenchymal components—fibroblasts in particular—are broadly implicated in maintenance of normal tissue structure or architecture as well as support of tissue homeostasis via production of soluble factors, basement membrane, and ECM components. Perturbation of fibroblast phenotypes to an activated state is an anatomically conserved feature of many solid cancers and some inflammatory conditions (25), and while activated fibroblasts in disease states generally express ECM components and immune-modulatory factors, granular features of fibroblast activation programs are tissue- and disease-specific. Though activated fibroblasts in cancer carry out diverse functions to promote tumor progression, normal fibroblasts serve to restrain tumor formation in promoting the ordered tissue structure that must be overcome to enable cancer formation or progression. We propose KITL as a tumor-restraining stromal mechanism in the pancreas, raising the possibility that specific effectors downstream of KITL signaling may hold utility for cancer progression. Future efforts will aim to investigate the significance of KITL signaling in the specific context of low-grade PanIN lesions, as these are the lesions found in adult human pancreas (19, 20).

While our study was restricted to the pancreas, these findings fit within a broader context of prior studies implicating mesenchymal KITL and/or LEPR-positive mesenchyme as critical regulators of tissue homeostasis and normal tissue function in diverse organ sites. As briefly discussed above, LEPR-positive mesenchymal cells in the bone marrow associate tightly with endothelial cells and form a niche critical for hematopoietic stem cells (22). Interestingly, upon tissue damage such as irradiation or chemotherapy requiring regeneration of hematopoietic stem cells, LEPR-positive mesenchymal cells differentiate into adipocytes which in turn produce KITL to enable a functional niche and support hematopoietic regeneration (30). Complementary mesenchymal and signaling components were recently shown to support normal tissue homeostasis and suppress inflammation in brown adipose tissue (BAT): LEPR-positive mesenchyme supports adaptive thermogenesis and restrains inflammation in BAT (23), while endothelial cell-derived KITL/SCF signals to KIT on brown adipocytes to promote homeostatic lipid accumulation when thermogenesis is inhibited (31). As the stellate cells under investigation in our study are also lipid-storing cells, these studies raise the possibility that lipid-storing stromal cells engage KITL signaling to promote tissue homeostasis and limit inflammation more broadly across organs.

## Methods

### Human tissue samples

All experiments with human patient-derived material were performed with approval of the Oregon Health & Science University and Memorial Sloan Kettering Cancer Center Institutional Review Boards. Sections from formalin-fixed, paraffin-embedded human PDAC patient tissue samples harboring benign adjacent pancreas tissue were donated to the Oregon Pancreas Tissue Registry program with informed written patient consent in accordance with full approval by the OHSU Institutional Review Board, or were obtained with informed consent of biospecimen collection with full approval by the MSKCC Institutional Review Board.

### Animals

All experiments involving mice were reviewed and overseen by the Institutional Animal Care and Use Committees at OHSU and MSKCC in accordance with National Institutes of Health guidelines for the humane treatment of animals. Male and female mice were used for all experiments, with ages specified in the experimental sections to follow. Littermate controls were used whenever possible. Animals included in pancreatitis and PDAC experiments were assessed daily based on score sheets with criteria including body condition scoring and physical examination to ensure humane treatment. Orthotopic tumors were grown to a maximum diameter of 1.0 cm based on institutional guidelines. Maximal burden was not exceeded with any animal. The following mice were used in this study, all purchased from the Jackson Laboratory: C57BL/6J (000664), *Rosa26^mTmG^* (007676), *Fabp4-Cre* (005069), *Kitl^flox^* (017861), *Trp53^frt^* (017767), *Kras^FSF-G12D^* (023590). The *Pdx1-FlpO* mouse strain was kindly provided by Dr. Michael Ostrowski (Medical University of South Carolina).

### Pancreatitis induction

Acute pancreatitis was induced in male and female mice at 8 weeks of age by intraperitoneal injection of caerulein (80 µg/kg, Sigma-Aldrich C9026) 8 times per day with 1 h between injections, on 2 consecutive days. Mice were then euthanized 2 days after the final caerulein injection and pancreata were collected.

### Orthotopic transplantation of PDAC cells

The 6419c5 and FC1245 cell lines were derived from autochthonous PDAC in the Kras^LSL-G12D/+^;Trp53^LSL-R172H/+^;Pdx1-Cre genetically engineered mouse model of pure C57BL/6J background, and were kindly provided by Dr. Ben Stanger (University of Pennsylvania) and Dr. David Tuveson (Cold Spring Harbor Laboratory), respectively. Male or female mice at 8-10 weeks of age were anesthetized and orthotopically implanted with 5 x 10^4^ (6419c5) or 5 x 10^3^ (FC1245) PDAC cells in a 50% Matrigel solution into the body of the pancreas. Tumor progression was monitored longitudinally by high-resolution ultrasound using the Vevo 2100 imaging system. Mice were euthanized and tumors collected either when the first mouse of the experiment reached humane endpoint, or at different time points when each individual mouse in the experiment reached humane endpoint.

### Single-cell RNA-seq

#### Cell isolation

To isolate healthy PSCs, pancreata were harvested from *Rosa26^mTmG/+^;Fabp4-Cre* mice at 6-9 weeks of age, trimmed to remove any associated adipose tissue, minced with scissors, digested with 0.02% Pronase (Sigma-Aldrich), 0.05% Collagenase P (Sigma-Aldrich), and 0.1% DNase I (Sigma-Aldrich) in Gey’s balanced salt solution (GBSS; Sigma-Aldrich) at 37°C for 10 minutes. Pancreata were further mechanically dissociated via serological pipette before returning to chemical dissociation at 37°C for 5 minutes. The resulting cell suspension was filtered through a 100 μm cell strainer nylon mesh. Cells were washed with GBSS, pelleted, and subject to red blood cell lysis via ACK lysis buffer (Thermo Fisher Scientific) for 3 minutes at room temperature. Then, cells were washed in cold FACS buffer (PBS containing 2% FBS), pelleted, and resuspended in FACS buffer. Cells were kept on ice as a single-cell suspension, then GFP-positive cells were isolated by FACS using a BD FACSAria III or BD FACSymphony S6.

To isolate CAFs, 8-week-old *Rosa26^mTmG/+^;Fabp4-Cre* mice were orthotopically implanted with FC1245 PDAC cells as described above. At 21 days post-implantation, pancreata were harvested, and any apparent normal pancreas tissue was trimmed away from the PDAC specimen. Tumors were briefly minced, placed in digestion media (DMEM with 1 mg/ml Collagenase IV, 0.1% soybean trypsin inhibitor, 50 U/ml DNase, and 0.125 mg/ml Dispase), and incubated at 37°C for 1 h. Whole tissue digests were centrifuged at 450 g for 5 min, then resuspended in 10 ml pre-warmed 0.25% Trypsin and incubated at 37°C for 10 min. Cold DMEM (10 ml) was added to the suspension, which was then passed through a 100 μm cell strainer. Cells were centrifuged as above, washed with DMEM containing 10% FBS and centrifuged again, then centrifuged as above and resuspended in 1 ml ACK red cell lysis buffer. Cells were incubated at room temperature for 3 minutes, then 9 ml FACS buffer added and cells centrifuged as above. Pelleted cells were counted and resuspended at 1 x 10^7^ cells/ml in FACS buffer, CD16/CD32 Fc block (BD 553141) added 1:20 and incubated at room temperature for 2 min, then biotinylated PDPN antibody (BioLegend 127404) was added 1:200. Cell suspensions were incubated on ice for 30 min. Cold FACS buffer was added, cells centrifuged at 300 g for 5 min at 4°C, and cell pellets were resuspended in 500 µl cold FACS buffer containing 1:1000 APC-streptavidin (BD 554067) and incubated for 30 min on ice protected from light. Cold FACS buffer (2 ml) was added, cells were pelleted as above and resuspended in cold FACS buffer containing SYTOX Blue Dead Cell Stain (Invitrogen S34857). Cells were incubated for 30 min on ice, washed with FACS buffer, pelleted, and resuspended in cold FACS buffer. GFP-positive PDPN-positive cells were isolated by FACS using a BD FACSAria III or BD FACSymphony S6.

#### Sequencing and analysis

The isolated pancreatic mesenchymal cells were immediately used for single-cell RNA-seq library preparation. Single cell capture and cDNA library generation were performed using the 10x Genomics Chromium single-cell 3’ library construction kit v2 (120267) according to the manufacturer’s instructions. Libraries were pooled prior to sequencing based on estimated cell number in each library per flow cytometry cell counts. Sequencing was performed on the Illumina NovaSeq 6000 platform at the OHSU Massively Parallel Sequencing Shared Resource, sequencing 20,000 read pairs per cell.

We aligned the sequenced reads to the mm10 mouse reference genome, and the unique molecule identifier (UMIs) for each gene in each cell were counted using the Cell Ranger (10x Genomics). Then, we imported the resulting gene expression matrices into R (version 4.0.3) and analyzed the data using the Seurat (32) pipeline (version 4.0.1). Genes had to be expressed in at least three cells to be considered for downstream analyses. Cells were filtered to retain those that contained at least 1,000 minimum unique genes expressed, no more than 5,000 unique genes, more than 200 total UMIs, and less than 10% of counts mapped to the mitochondrial genome. Batch correction was performed to integrate the samples from different conditions using the reciprocal PCA (RPCA) integration workflow (33) within Seurat. The first 30 principal components were selected for downstream analysis, based on the elbow point on the plot of standard deviations of principal components. UMAP was generated using the RunUMAP function with the same first 30 principal components used in clustering analysis.

We performed pseudotime trajectory analysis to elucidate the differentiation pathways of normal pancreatic and cancerous cells using Monocle 3 (v.1.3.7) (34). To achieve this, we first integrated our single-cell RNA-seq datasets using Harmony (v.0.1.1) (35) to correct for batch effects, enabling a unified visualization of cellular heterogeneity across samples. Subsequent trajectory inference with Monocle 3 was conducted using default parameters to order cells in pseudotime, thus highlighting the dynamic progression of cellular states. To visualize gene expression patterns along the trajectories, we utilized the ‘plot_cell_trajectory’ function, focusing on the expression of Kitl in the harmony-adjusted dimensional space.

### Mouse pancreatic stellate cells (mPSCs) isolation

Primary mPSCs were isolated from wild-type C57BL/6J (000664) mice from The Jackson Laboratory at 8-9 weeks of age. Our isolation protocol is adapted from previously described methods (36, 37) with some minor modifications. Healthy pancreatic tissues from eight male mice were pooled, trimmed, and digested in Hank’s balanced salt solution (HBSS; Sigma Aldrich, H8264) containing 0.5 mM of magnesium chloride hexahydrate (MgCl_2_ x 6H_2_O; Sigma Aldrich, M9272), 10 mM HEPES (Cytiva, SH30237.01), 0.13% Collagenase P (Roche, 11213873001), 0.1% protease (Sigma Aldrich, p5417), and 0.001% DNase (Roche, 04716728001) for 7 minutes in shaking water bath (120 cycles/min) at 37℃. Remaining connective and adipose tissues were removed before the second incubation at 37℃ in a shaking water bath (80 cycles/min) for an additional 7 minutes. Digested tissues were then filtered through a 250 µm nylon mesh (Thermo Fisher Scientific, 87791) and centrifuged at 450 g for 10 minutes at 4℃. The cell pellet was washed in Gey’s balanced salt solution (GBSS) containing 120 mM salt (NaCl; Sigma Aldrich, S3014) and 0.3% BSA (Fisher Scientific, BP9703100) before repeating the centrifugation step above. Upon removing the wash buffer, cells were resuspended in GBSS + NaCl containing 0.3% BSA, to which equal volume of 28.7% solution of Nycodenz (ProteoGenix, 1002424) in GBSS - NaCl were added and mixed well. The cell suspension in Nycodenz is then gently layered beneath GBSS containing 120 mM NaCl and 0.3% BSA using a long needle and subjected to centrifugation at 1400xg for 20 minutes at 4℃. Primary quiescent PSCs were carefully harvested from the interface using sterile pipette and washed with GBSS + NaCl containing 0.3% BSA. Cells were pelleted and plated into multiple wells of a 6-well dish in Iscove’s modified Dulbecco’s medium (IMDM; Cytiva, SH30228.02) containing 10% FBS (VWR, 97068-085) and 1% Antibiotics-Antimycotic (Thermo Fisher Scientific, 15240-062). Cell culture was maintained in a humidified atmosphere at 37℃ with 5% CO_2_.

### Stable *Kitl* knock down and overexpression in pancreatic stellate cells

*Kitl* knock down (sh*Kitl*) and overexpression (*Kitl* OE) mPSC-1 cell lines were generated using Mission Lentiviral shRNA (Millipore Sigma, Clone ID: TRCN0000067872) and *Kitl* open reading frame lentivirus (Genecopoeia, EX-Mm03868-Lv158) respectively. Vector PLKO.1 Neo (sh*Ctrl*; Addgene, 13425) and *Egfp* open reading frame (*Egfp* OE; Genecopoeia, EX-EGFP-Lv158) were included as controls. Immortalized mPSC-1 cells were transduced with specified lentiviral particles for 48 hours prior to selection with 1 mg/mL Geneticin (Fisher Scientific, 10131035) for 4 days. KITL protein and transcript expression were then quantified using qPCR and ELISA to assess silencing and overexpression efficiency. Stable cells were maintained in a humidified atmosphere at 37℃ with 5% CO_2_ and routinely passed in DMEM (Thermo Fisher Scientific, 11965118) containing 10% FBS (VWR, 97068-085) 1 mM sodium pyruvate (Thermo Fisher Scientific, 11360070), and 1% Antibiotics-Antimycotic (Thermo Fisher Scientific, 15240-062).

### RNA-sequencing of sh*Kitl* and *Kitl* ORF pancreatic stellate cells

Total RNA was isolated using RNeasy Microkit (Qiagen, 74004) per manufacturer’s instructions and quantified using NanoDrop microvolume spectrophotometer before submission for bulk RNA-sequencing. RNA library preparation, sequencing, and analysis were conducted at Azenta Life Sciences (South Plainfield, NJ, USA) as follows. Total RNA samples were quantified using Qubit 2.0 Fluorometer (Life Technologies, Carlsbad, CA, USA) and RNA integrity was checked using Agilent TapeStation 4200 (Agilent Technologies, Palo Alto, CA, USA). ERCC RNA Spike-In Mix (Cat: #4456740) from ThermoFisher Scientific, was added to normalized total RNA prior to library preparation following manufacturer’s protocol. Total RNA underwent polyA selection and RNA sequencing libraries preparation using the NEBNext Ultra II RNA Library Prep Kit for Illumina using manufacturer’s instructions (NEB, Ipswich, MA, USA). Briefly, mRNAs were initially enriched with Oligod(T) beads. Enriched mRNAs were fragmented for 15 minutes at 94 °C. First strand and second strand cDNA were subsequently synthesized. cDNA fragments were end repaired and adenylated at 3’ends, and universal adapters were ligated to cDNA fragments, followed by index addition and library enrichment by PCR with limited cycles. The sequencing library was validated on the Agilent TapeStation (Agilent Technologies, Palo Alto, CA, USA), and quantified by using Qubit 2.0 Fluorometer (Invitrogen, Carlsbad, CA) as well as by quantitative PCR (KAPA Biosystems, Wilmington, MA, USA). The sequencing libraries were multiplexed and clustered onto a flowcell on the Illumina NovaSeq instrument according to manufacturer’s instructions. The samples were sequenced using a 2x150bp Paired End (PE) configuration at an average of 30 million reads per sample. Image analysis and base calling were conducted by the NovaSeq Control Software (NCS). Raw sequence data (.bcl files) generated from Illumina NovaSeq was converted into fastq files and de-multiplexed using Illumina bcl2fastq 2.20 software. One mismatch was allowed for index sequence identification.

After investigating the quality of the raw data, sequence reads were trimmed to remove possible adapter sequences and nucleotides with poor quality. The trimmed reads were mapped to the reference genome GRCm38.91 (mm10) available on ENSEMBL using the STAR aligner v.2.5.2b. The STAR aligner is a splice aligner that detects splice junctions and incorporates them to help align the entire read sequences. BAM files were generated as a result of this step. Unique gene hit counts were calculated by using feature Counts from the Subread package v.1.5.2. Only unique reads that fell within exon regions were counted. The gene hit counts table was used for downstream differential expression analysis. Using DESeq2, a comparison of gene expression between the groups of samples was performed. The Wald test was used to generate p-values and Log2 fold changes. Genes with adjusted p-values < 0.05 and absolute log2 fold changes > 1 were called as differentially expressed genes for each comparison. Volcano plot visualization of significant DEGs were performed in Galaxy (38) using the ggplot2 R package. Significant gene labels from top gene ontologies categories were included.

Functional enrichment analysis was performed using enrichR (39) on the statistically significant set of genes by implementing Fisher exact test (GeneSCF v1.1-p2). Significance of tests was assessed using adjusted p-values defined by enrichR. Enrichment bar plots were generated using srPlot (40) to include Top 10 upregulated and downregulated gene ontology categories.

### Immunohistochemistry, immunofluorescence, and lipid staining

#### Mouse and human tissue sample staining

Standard protocols were performed for IHC. Briefly, tissue samples were fixed overnight in 10% neutral-buffered formalin (Sigma-Aldrich, HT501128-4L) and submitted to MSKCC Laboratory of Comparative Pathology or Molecular Cytology Core Facility for paraffin embedding, sectioning and H&E sectioning. Sectioned tissues were deparaffinized using CitriSolv (Fisher Scientific, 22-143-975) and rehydrated in ethanol series (Decon labs, 2701) before undergoing antigen retrieval using citrate or tris based antigen unmasking solution (Vector laboratories, H3300, H3301). The slides were then blocked with 8% BSA (Fisher Bioreagents, BP9703100) for 1 hour at room temperature and incubated in primary antibodies at 4℃ overnight. Primary antibodies for ɑSMA (Cell Signaling Technology, 19245S), PDPN (eBio 8.1.1 Invitrogen, 14538182), GFP (Thermo Fisher, A10262; Abcam, ab1218; Rockland Immunochemicals, 600-101-215), pan-cytokeratin (Thermo Fisher Scientific, MA5-13156), CD31 (R&D AF3628 or Abcam ab7388), biotinylated anti-c-Kit (R&D BAF1356), Vimentin (Cell Signaling Technology 5741 D21H3 XP), or pancreatic amylase (Thermo Scientific PA5-25330) were diluted at 1:200-1:400 in 8% BSA in PBS. The next day, slides were washed with PBS (Biotum, 22020) and incubated in α-chicken Alexa Fluor 488 (Thermo Fisher Scientific, A32931), α-rabbit Alexa Fluor 647 (Fisher Scientific, A21245), α-Syrian hamster Alexa Fluor 647 (Abcam, ab180117), or α-mouse Alexa Fluor 555 (Fisher Scientific, A21424) secondary antibodies at 1:200-1:400 dilution for 1 hour at room temperature. Tissue slides were washed with PBS and mounted with Vectashield mounting media containing DAPI (Vector laboratories, H-1200-10).

All images were acquired on a Carl Zeiss LSM880 laser-scanning confocal inverted microscope using 20x, 40X, or 63X objective. Whole slide scans were completed by MSKCC Molecular Cytology Core Facility. Image analysis was performed using QuPath quantitative pathology and FIJI/ImageJ open source software. Where applicable, co-localization analysis was performed using the JaCop plugin in ImageJ.

#### Cell staining and imaging

Cells seeded in chamber slides were fixed in 4% paraformaldehyde for 15 minutes and permeabilized with 0.1% Triton X-100 for 10 minutes before undergoing blocking in 5% BSA for 1 hour at room temperature. Sample slides were then probed with ɑSMA primary antibodies (ThermoFisher, MA5-11547) overnight at 4℃ followed by standard Alexa Fluor 647-conjugated secondary antibody (Fisher Scientific, A21235) incubation for an hour at room temperature. Upon repeating standard washing steps, slides were mounted for imaging using Vectashield mounting media containing DAPI (Vector laboratories, H-1200-10). For lipid staining, cells seeded in chamber slides were stained with Nile Red (MCE, HY-D0718) at 1 µM final working concentration for 10 minutes and counterstained with DAPI (Thermo Fisher Scientific, 62248). Nile Red signals were detected at excitation/emission wavelengths 559 nm/ 635 nm.

### Two-plex fluorescence *in situ* hybridization

Transcript expression on tissues, except where RNAscope was indicated, was performed using the Thermo Fisher Scientific ViewRNA ISH Tissue Assay kit (two plex) for use on mouse and human tissue samples. Briefly, samples were first permeabilized with controlled protease digestion, followed by incubation with proprietary probe-containing solution, according to the manufacturer’s instructions. During incubation, samples had to remain fully submerged. After hybridization with the probe, samples were washed, followed by sequential hybridization with the preamplifier and amplifier DNA. In accordance with the manufacturer’s instructions, hybridizations were performed with the preamplifier, amplifier and fluorophore. Mounting medium with DAPI (Vectashield Hardset mounting media with DAPI) was used to mount samples.

### RNAscope combined with immunohistochemistry

Paraffin-embedded tissue sections were cut at 5 μm and kept at 4°C. Samples were loaded into Leica Bond RX, baked for 30 mins. at 60°C, dewaxed with Bond Dewax Solution (Leica, AR9222), and pretreated with EDTA-based epitope retrieval ER2 solution (Leica, AR9640) for 15 mins. at 95°C. The probe mKitL (Advanced Cell Diagnostics, ready to use, no dilution, 423408) was hybridized for 2hrs. at 42°C. Mouse PPIB (ACD, Cat# 313918) and dapB (ACD, Cat# 312038) probes were used as positive and negative controls, respectively. The hybridized probes were detected using RNAscope 2.5 LS Reagent Kit – Brown (ACD, Cat# 322100) according to manufacturer’s instructions with some modifications (DAB application was omitted and replaced with Fluorescent CF594/Tyramide (Biotium,92174) for 20 mins. at RT).

After the run was finished, slides were washed in PBS and incubated in 5 μg/ml 4’,6-diamidino-2-phenylindole (DAPI) (Sigma Aldrich) in PBS for 5 min, rinsed in PBS, and mounted in Mowiol 4– 88 (Calbiochem). Slides were kept overnight at -20°C before imaging.

After the slides were scanned, the coverslips were removed and slides were loaded into Leica Bond RX for double IF staining. Samples were pretreated with EDTA-based epitope retrieval ER2 solution (Leica, AR9640) for 20 mins. at 100°C. The double antibody staining and detection were conducted sequentially. The primary antibodies against GFP (2ug/ml, chicken, abcam, ab13970) and CD31 (CD31/A647 (0.08, rb, abcam, ab182981) were incubated for 1h at RT. For rabbit antibodies, Leica Bond Polymer anti-rabbit HRP (included in Polymer Refine Detection Kit (Leica, DS9800) was used, for the chicken antibody, a rabbit anti-chicken (Jackson ImmunoResearch303-006-003) secondary antibodies were used as linkers for 8 min before the application of the Leica Bond Polymer anti-rabbit HRP for 8 min at RT. The Leica Bond Polymer anti-rabbit HRP secondary antibody was applied followed by Alexa Fluor tyramide signal amplification reagents (Life Technologies, B40953 and B40958) were used for IF detection. After the run was finished, slides were washed in PBS and mounted in Mowiol 4–88 (Calbiochem). Slides were kept overnight at -20°C before imaging.

### CODEX

#### Antibody panel development, CODEX staining and imaging

To construct an antibody panel visualizing pancreatic architecture in FFPE mouse samples using CODEX (29), conventional IHC staining was performed to screen for antibodies binding canonical markers of pancreatic epithelial cells [E-cadherin, Novus Biologicals #NBP2-33006 clone 1A4(asm-1); Amylase, Cell Signaling Technology #3796 clone D55H10], endothelial cells (CD31, Cell Signaling Technology #14472 clone 4A2), stromal cells (Vimentin, Cell Signaling Technology #70257 clone D3F8Q; α-SMA, Cell Signaling Technology #77699 clone D8V9E), leukocytes (CD45, Cell Signaling Technology #46173 clone D21H3) and lineage reporter (GFP, Rockland Immunochemicals #600-101-215 polyclonal). Identified antibody clones were then conjugated with oligonucleotide barcodes using Antibody Conjugation Kit (Akoya Biosciences). Prior to CODEX imaging, each conjugated antibody was validated following manufacturer instructions and tissue staining patterning was confirmed with published literature.

CODEX staining and imaging was performed as described in user manual (https://www.akoyabio.com/wp-content/uploads/2021/01/CODEX-User-Manual.pdf). In brief, 5 µm FFPE pancreas sections were mounted onto 22 mm x 22 mm glass coverslips (Electron Microscopy Sciences) coated in 0.1% poly-L-lysine (Sigma) and stained with using CODEX Staining Kit (Akoya Biosciences). A cocktail of above-conjugated antibodies were incubated with tissue overnight at 4°C. On the next day, fluorescent oligonucleotide-conjugated reporters were combined with Nuclear Stain and CODEX Assay Reagent (Akoya Biosciences) in sealed light-protected 96-well plates (Akoya Biosciences). Automated fluidics exchange and image acquisition were performed using the Akoya CODEX instrument integrated with a BZ-X810 epifluorescence microscope (Keyence) and CODEX Instrument Manager (CIM) v1.30 software (Akoya Biosciences). The exposure times were as follows: E-cadherin, barcode BX006, 600 ms; Amylase, barcode BX031, 250 ms; Vimentin, barcode BX025, 300 ms; α-SMA, barcode BX052, 250 ms; CD31, barcode BX002, 350 ms; CD45, barcode BX007, 400 ms; GFP, barcode BX041, 250 ms. All images were acquired using a CFI plan Apo I 20×/0.75 objective (Nikon). “High resolution” mode was specified in Keyence software to reach a final resolution of 377.44 nm/pixel.

#### Processing of CODEX images and analysis

Image stitching, drift compensation, deconvolution, z-plane selection, and background subtraction were performed using the CODEX processor v1.7 (Akoya Biosciences) per manufacture instruction (https://help.codex.bio/codex/processor/technical-notes). Individual channel images were then imported into ImageJ v1.53t for analyses as described below.

Total pancreatic areas were annotated by sum of Amylase^+^ and Ecadherin^+^ region. Immune cells were defined by DAPI and CD45 double positivity while vasculature area was annotated by CD31+ region. Vimentin and α-SMA signal were used to mark total and activated fibroblast cells, respectively. GFP positivity was used to track PSC lineage-derived cells.

### Flow cytometry

To analyze c-KIT expression, normal pancreas tissues were harvested from wild-type C57BL/6J mice aged 6-9 weeks and digested as described above. Following ACK lysis, cells were incubated with CD16/CD32 antibody (BD Biosciences, 553141) to block Fc receptors for 2 minutes at room temperature. Cells were then stained with the following for 30 minutes on ice: SYTOX Blue Dead Cell Stain (Invitrogen S34857); biotinylated m-SCF R/c-KIT antibody (R&D Systems BAF1356). Cells were then washed with cold FACS buffer, pelleted, then stained with PE/Cy7 Streptavidin (Biolegend, 405206), anti-mouse CD31 APC (Invitrogen 17-0311-82), anti-mouse EpCAM (CD326) FITC (Invitrogen 11-5791-82) for 30 minutes on ice, before cells were washed with FACS buffer, pelleted, then resuspended in cold FACS buffer for flow cytometry.

To analyze epithelial cells, immune cells, and c-KIT, pancreata from C57Bl/6J mice aged 6-9 weeks old were harvested and digested as described above. After ACK lysis, cells were incubated with CD16/CD32 antibody (BD Biosciences 553141) for 2 minutes at room temperature. Cells were then stained with SYTOX Blue and a biotinylated c-KIT antibody on ice for 30 minutes on ice, were washed with cold FACS buffer, pelleted, then stained with PE/Cy7 Streptavidin (Biolegend 405206), anti-mouse CD45 PE-Cyanine 5 (Invitrogen 15-0451-82), anti-mouse EpCAM (CD326) FITC (Invitrogen 11-5791-82) for 30 minutes on ice. Cells were washed with FACS buffer, pelleted, then resuspended in cold FACS buffer for flow cytometry.

### Gene expression analysis by qPCR

Total RNA was isolated using RNeasy Microkit (Qiagen, 74004) per manufacturer’s instructions and quantified using NanoDrop microvolume spectrophotometer. 500 ng to 1 µg of RNA was reverse transcribed using iScript reverse transcriptase supermix (Bio-Rad, 1708841) to produce cDNA. Real-time PCR was performed using Power SYBR Green PCR master mix (Thermo Fisher, 4367659). Gene specific primer pairs were designed using the NCBI Nucleotide database or acquired from Millipore Sigma. Gene expression is normalized to reference gene *Rplp0*. Primer pair sequences were as follows: *Rplp0* Forward 5’-GTGCTGATGGGCAAGAAC-3’ Reverse 5’-AGGTCCTCCTTGGTGAAC-3’, m*Kitl* Forward 5’-TTATGTTACCCCCTGTTGCAG-3’ Reverse 5’-CTGCCCTTGTAAGACTTGACTG-3’, m*Kit* Forward 5’-GAGACGTGACTCCTGCCATC-3’ Reverse 5’-TCATTCCTGATGTCTCTGGC-3’, m*Acta2* Forward 5’-AGCCATCTTTCATTGGGATGGA-3’ Reverse 5’-CATGGTGGTACCCCCTGACA-3’.

### ELISA quantikine assay

Immortalized parental and sh*Kitl* mPSC-1 cells were seeded into 6 well dish at 3 x 10^5^ confluency in growth media containing DMEM (Thermo Scientific, 11965126), 10% VWR Seradigm FBS (VWR, 97068-085), 1 mM Sodium Pyruvate (Thermo Scientific, 11360070), and 1% Antibiotics-Antimycotic (Thermo Fisher Scientific, 15240-062). Primary PSCs were seeded in 6 well dish at 1 x 10^4^ confluency in Iscove’s modified Dulbecco’s medium (IMDM; Cytiva, SH30228.02) containing 10% FBS (VWR, 97068-085) and 1% Antibiotics-Antimycotic (Thermo Fisher Scientific, 15240-062). Conditioned media were collected at indicated time points and concentrated using Vivaspin Turbo 20 3K MWCO concentrator (Cytiva, 28932358) in accordance with manufacturer’s protocol. Concentrated supernatants were quantified using Pierce BCA Protein Assay Kit (Thermo Fisher, 23225) and mouse KITL protein quantification was performed using Mouse SCF Quantifikine ELISA Kit (R&D Systems, MCK00) according to manufacturer’s protocol.

### Statistical analysis

No statistical methods were used to predetermine sample sizes. The experiments were not randomized. For animal studies, a minimal number of mice was selected based on preliminary studies, with an effort to achieve a minimum of n = 3, mostly n = 5-10 mice per treatment group for each experiment. Age-matched mice were selected for experiments. For histological staining quantification, analyses were performed in a blinded fashion. For batch-processed images, image analyses were done in an unbiased manner using image analysis software. Some western blots and RT-qPCR assays were performed by a researcher blind to the experimental hypothesis. Animals were excluded if an animal needed to be removed from an experiment early for reasons seemingly unrelated to tumor burden. All experiments were performed and reliably reproduced at least two independent times. GraphPad Prism 9 was used to generate graphs and for statistical analyses. Groups were tested for normality. Statistical significance was calculcated for two unmatched groups by unpaired *t*-test with Welch’s correction or Mann-Whitney test. One- or two-way ANOVAs were used for more than two groups as specified, followed by Tukey’s multiple comparisons tests. Datasets are presented as mean ± s.e.m. *P* values under 0.05 were considered significant. Data distribution was assumed to be normal, but this was not formally tested.

## Acknowledgements

We thank Mark Berry, Wenfei Kang, and all members of the Sherman lab for helpful discussion of this work. This study was supported by NIH grants R01 CA229580 and R01 CA250917 (to M.H.S.), NIH grant P01 CA244114 (to M.H.S. and Y.H.), NIH grant R01 GM147365 (to Z.X.), and a Silver Family Innovation Fund Award (to Z.X.). We thank members of the OHSU Histopathology Shared Resource, Advanced Light Microscopy Shared Resource, Flow Cytometry Shared Resource, and Massively Parallel Sequencing Shared Resource for supporting this study, with financial support from NIH-NIH Cancer Center Support Grant P30 CA069533. We also thank members of the MSKCC Flow Cytometry Core, Molecular Cytology Core, and Center of Comparative Medicine & Pathology for supporting this work, with financial support from NIH-NCI Cancer Center Support Grant P30 CA008748.

## Data availability statement

The data generated in this study will be made publicly available in the Gene Expression Omnibus prior to publication.

---

**Supplementary Figure S1:**
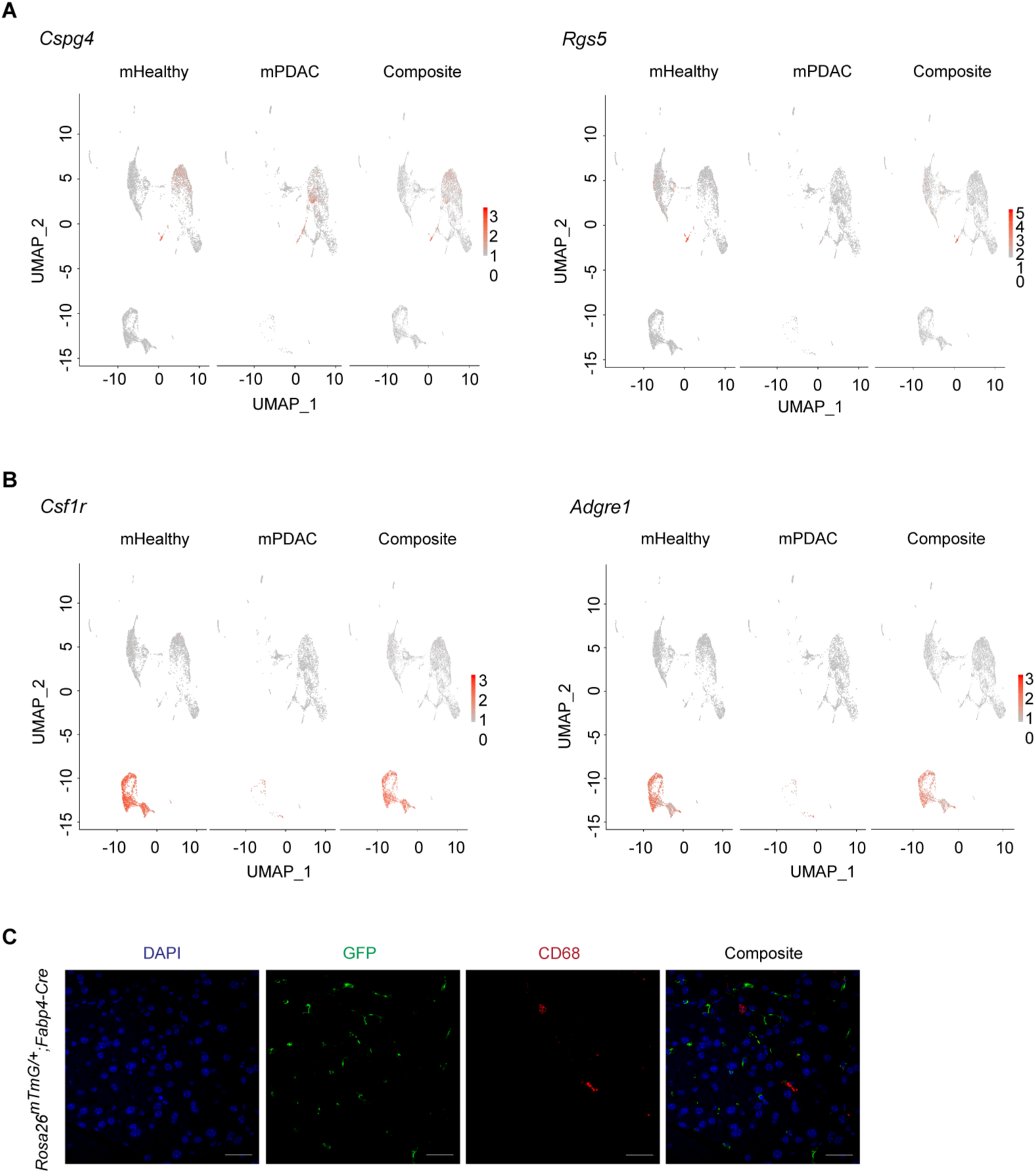
scRNA-seq reveals gene expression programs in PSCs and PSC-derived CAFs. **A,** UMAP (uniform manifold approximation and projection) visualization of *Cspg4* and *Rgs5* gene expression in scRNA-seq dataset of pancreatic stellate cells (PSCs) and PSC-derived cancer associated fibroblasts (CAFs) isolated from healthy pancreas and orthotopic tumors respectively. **B,** UMAP (uniform manifold approximation and projection) visualization of *Csf1r* and *Adgre1* gene expression in pancreatic stellate cells (PSCs) and PSC-derived cancer associated fibroblasts scRNA-seq dataset (n = 2 replicates pooled from n = 5 mice per arm). **C,** Representative images of IHC staining for GFP (green) and CD68 (red) in normal pancreas tissue from *Rosa26^mTmG/+^;Fabp4-Cre* mice (n = 3 mice). Scale bar: 50 μm.

**Supplementary Figure S2:**
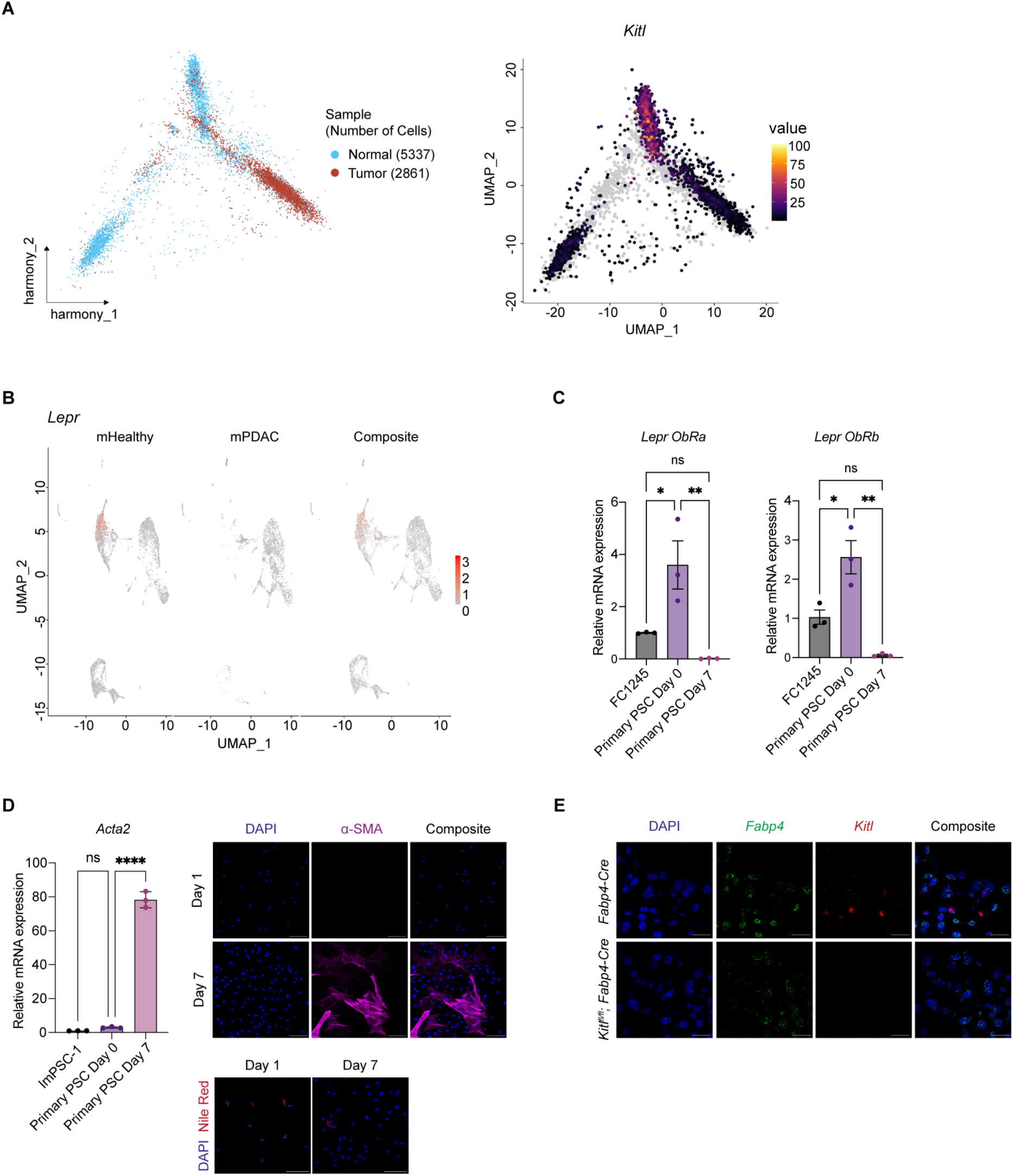
*Kitl* is expressed by healthy pancreatic mesenchyme and reduced upon activation to a CAF phenotype. **A,** (Left) UMAP illustrating the cellular landscape of normal pancreatic (blue) and PDAC (red) mesenchymal cells, comprising 5,337 normal and 2,861 tumor cells. Harmony was used to integrate the datasets and correct for batch effects. (Right) Monocle 3 trajectory analysis was used to depict expression of the *Kitl* gene along the inferred pseudotime trajectory. Cells are colored based on *Kitl* expression levels, with values ranging from low (black) to high (yellow), revealing the spatial and temporal expression patterns of *Kitl* (n = 2 replicates pooled from n = 5 mice per arm). **B,** UMAP (uniform manifold approximation and projection) visualization of *Lepr* gene expression in normal pancreatic stellate cells (PSCs) and PSC-derived cancer associated fibroblasts scRNA-seq dataset (n = 2 replicates pooled from n = 5 mice per arm). **C,** qRT-PCR of *Lepr ObRa* and *Lepr ObRb* isoforms in primary pancreatic stellate cells harvested at indicated time point. FC1245 PDAC cell line was included as a reference point. Data represents biological triplicates plotted as mean ± SEM. Significance was determined by ordinary one-way ANOVA; ns = not significant, *P≤ 0.05, **P≤ 0.01. **D,** Left: qRT-PCR analysis of *Acta2* in quiescent (Day 0) and activated (Day 7) primary pancreatic stellate cells (PSCs) with immortalized ImPSC-1 included as reference point. Data represents biological triplicate plotted as mean ± SEM. Significance was determined by ordinary one-way ANOVA; ns = not significant, ****P≤ 0.0001. Right: Representative immunofluorescence staining for ɑ-SMA and Nile Red staining in primary pancreatic stellate cells fixed at indicated time point (n= 3 biological replicates). Scale Bar: 100 μm. **E,** Representative images of RNA FISH staining for *Fabp4* (green) and *Kitl* (red) in murine normal pancreas between *Fabp4-Cre* control and *Kitl^fl/fl^;Fabp4-Cre* mice. Scale bar, 10 μm.

**Supplementary Figure S3:**
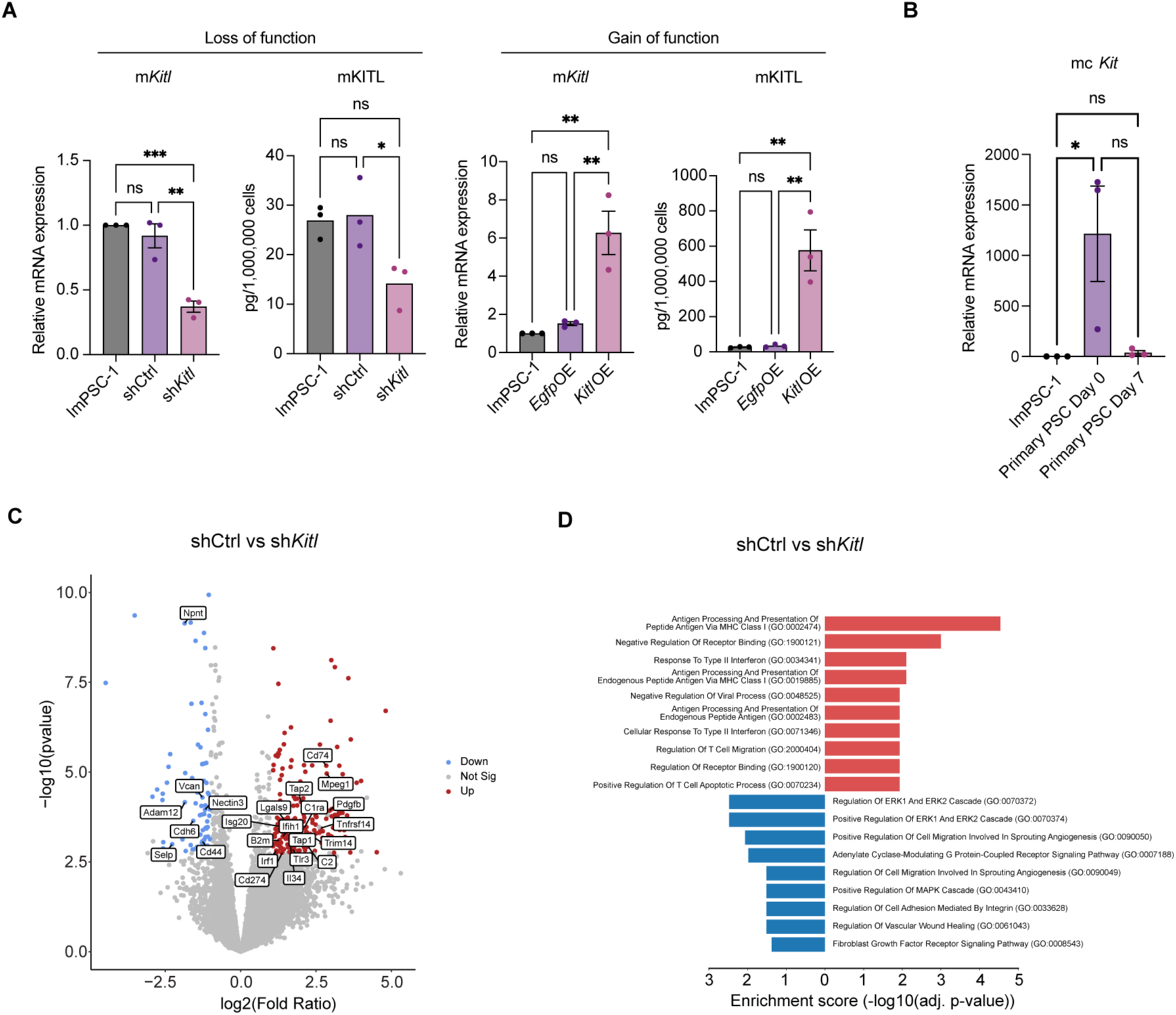
Stromal KITL promotes regulation of pancreas tissue architecture. **A,** Soluble *Kitl* transcript level (left) and protein secretion (right), quantitated using qRT-PCR and ELISA respectively, of immortalized pancreatic stellate cells (ImPSC-1) expressing stable *Kitl* knockdown (sh*Kitl*) or overexpression (*Kitl* OE). Parental ImPSC-1 serves as control for both stable cell lines. Data represents biological triplicates plotted as mean ± SEM. Significance was determined by ordinary one-way ANOVA; ns = not significant, *P≤ 0.05, **P≤ 0.01, ***P≤ 0.001. **B,** qRT-PCR of *cKit* in quiescent (Day 0) and activated (Day 7) primary pancreatic stellate cells (PSCs) with immortalized ImPSC-1 included as reference point. Data represents biological triplicate plotted as mean ± SEM. Significance was determined by ordinary one-way ANOVA; ns = not significant, *P≤ 0.05. **C**, Volcano plot of all upregulated, non-significant, and downregulated differentially expressed genes as defined by the Wald test (p.adj <0.05 and log_2_FC >1) from *Kitl* knock down (sh*Kitl*) ImPSC-1 bulk-RNA seq dataset with representative gene labels included. Data is representative of 3 biological repeats. **D,** Gene ontology (GO) analysis of upregulated and downregulated differentially expressed genes in immortalized pancreatic stellate cells (ImPSC-1) with *Kitl* stable knockdown. Top 10 enrichment categories ranked by adjusted p-values plotted in each direction.

**Supplementary Figure S4:**
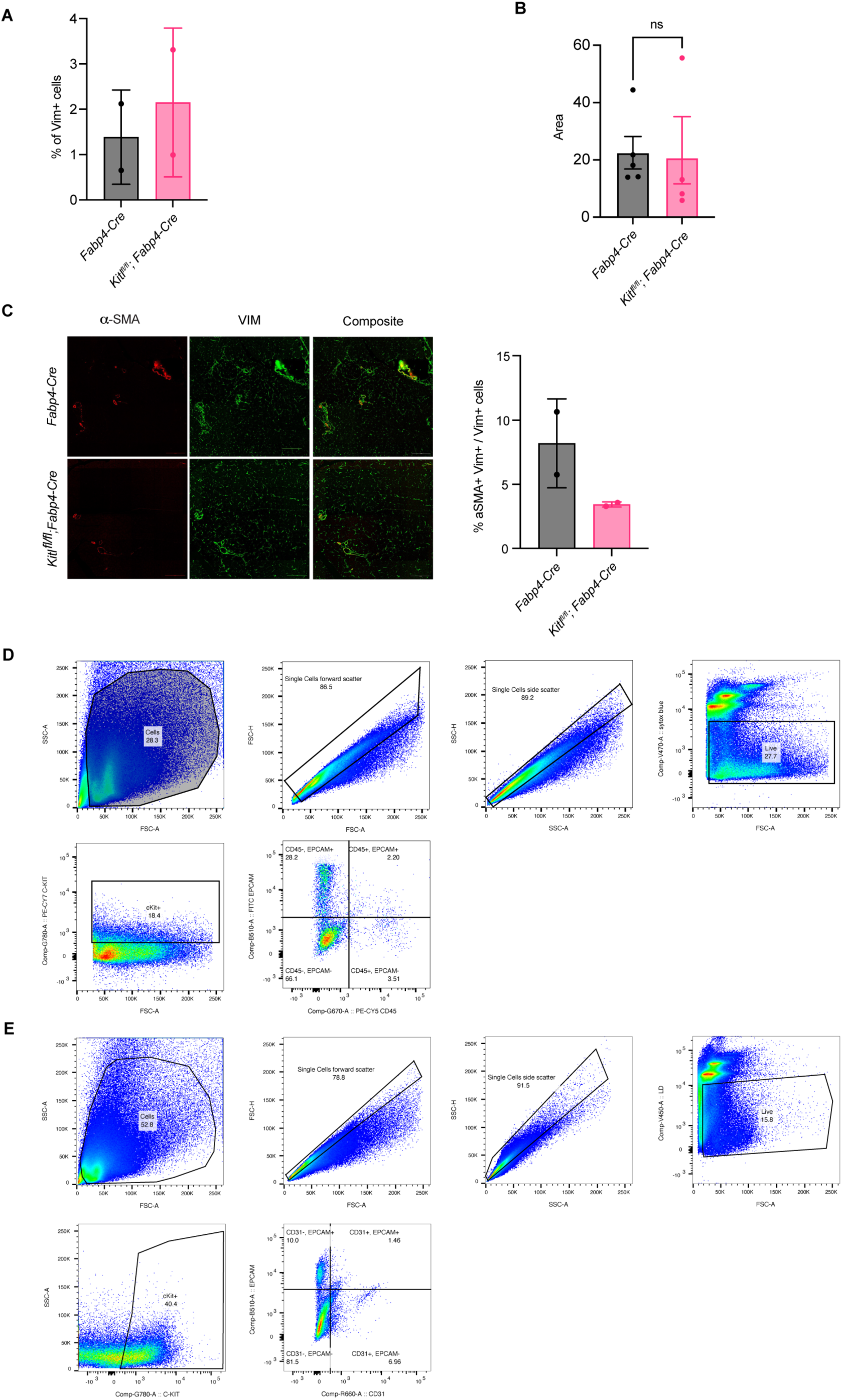
Stromal KITL promotes pancreas tissue homeostasis. **A,** CODEX quantification of Vimentin (VIM) between healthy pancreata of *Fabp4-Cre* control and *Kitl^fl/fl^;Fabp4-Cre* mice (n = 2 mice per arm). Data are represented as as mean ± SD. **B,** IHC quantification of CD31^+^ area between healthy pancreata of *Fabp4-Cre* control and *Kitl^fl/fl^;Fabp4-Cre* mice (n = 5 for control; n = 4 for *Kitl^fl/fl^;Fabp4-Cre*). Data are represented as mean ± SEM. **C,** Representative images of CODEX staining (left) and quantification (right) of αSMA and Vimentin (VIM) between healthy pancreata of *Fabp4-Cre* control and *Kitl^fl/fl^;Fabp4-Cre* mice. CODEX quantification (right) of VIM expression (n = 2 mice per arm). Data are represented as as mean ± SD. **D,** Flow cytometry analysis of cKIT receptor, EpCAM, and CD45 in healthy pancreata of C57BL/6J age-matched male mice (n = 5). **E,** Flow cytometry analysis of cKIT receptor, EpCAM, and CD31 in healthy pancreata of C57BL/6J age-matched male mice (n = 5). *, P < 0.0332; **, P < 0.0021; ***, P < 0.0002; ****, P < 0.0001 Mann-Whitney unpaired t test; ns, not significant.

## Notes

### Competing Interest Statement

The authors have declared no competing interest.

